# A gate-and-switch model for head orientation behaviors in *C. elegans*

**DOI:** 10.1101/291005

**Authors:** Marie-Hélène Ouellette, Melanie J. Desrochers, Ioana Gheta, Ryan Ramos, Michael Hendricks

## Abstract

The nervous system seamlessly integrates perception and action. This ability is essential for stable representation of and appropriate responses to the external environment. How the sensorimotor integration underlying this ability occurs at the level of individual neurons is of keen interest. In *Caenorhabditis elegans*, RIA interneurons receive input from sensory pathways and have reciprocal connections with head motor neurons. Through separate physiological mechanisms, RIA simultaneously encodes both head orientation and sensory stimuli. Based on these observations, we proposed a model for how RIA may integrate these two signals to detect the spatial distribution of stimuli across head sweeps and generate directional head responses. Here, we show that blocking synaptic release from RIA disrupts head orientation behaviors in response to unilaterally presented stimuli. We found that sensory encoding in RIA is gated according to head orientation. This dependence on head orientation is independent of motor encoding in RIA, suggesting a second, posture-dependent pathway upstream of RIA. This gating mechanism may allow RIA to selectively attend to stimuli that are asymmetric across head sweeps. Attractive odor removal during head bends triggers rapid head withdrawal in the opposite direction. Unlike sensory encoding, this directional response is dependent on motor inputs to and synaptic output from RIA. Together, these results suggest that RIA is part of a sensorimotor pathway that is dynamically regulated according to head orientation at two levels: the first is a gate that filters sensory representations in RIA, and the second is a switch that routes RIA synaptic output to dorsal or ventral head motor neurons.

## Introduction

Perception and action are intimately linked in the central nervous system. Our movements often have immediate sensory consequences, including changes in visual, proprioceptive, and mechanosensory sensations. Likewise, vocalization produces auditory self-stimuli, and inhalation can result in olfaction. To ensure accurate internal representations of the world, reafferent sensory stimulation caused by our own behavior must be processed and interpreted differently from other stimulus sources (Straka et al., 2018). In addition to reafference, many of our sensory systems predict the consequences of our actions in advance via inputs from motor areas, called efference copies or corollary discharge (Crapse and Sommer, 2008). Through processes that are only partially understood, our senses are precisely filtered and modulated in real time to compensate for ongoing behaviors and the sensory consequences of our actions. While some of the neural circuits that underlie these functions have been anatomically characterized to varying degrees, the sites and molecular mechanisms of this type of sensorimotor integration remain elusive in all but a few cases. The nematode *C. elegans*, which has a completely mapped and largely invariant nervous system, quantifiable behaviors, and a suite of genetic and imaging tools, provides an opportunity to probe these events at cellular resolution and directly link them to behavior (Calhoun and Murthy, 2017; White and Southgate, 1986).

The primary mode of *C. elegans* navigation is a biased random walk consisting of bouts of forward movement punctuated by quasi-random reorientations achieved through clustered sequences of reversals and turns. Directional movement is achieved by measuring changes in sensory input over time during forward bouts and modulating reorientation probability accordingly (Pierce-Shimomura et al., 1999). This behavior is directly analogous to the runs and tumbles of bacterial chemotaxis (Berg, 1975). *C. elegans* lie on their side, undulating in the dorsoventral plane as they crawl in a sinusoidal motion. Sensory structures located at the tip of the nose sweep back and forth as the head bends during forward movement. In contrast to the biased random walk, several navigation behaviors suggest that reafferent sensory input caused by head bending during locomotion is exploited as a way to sample the spatial distribution of environmental stimuli and guide navigation, a simple form of active sensing. For example, in radial temperature gradients, *C. elegans* will crawl in precisely curved trajectories to stay within a favored temperature range, a behavior called isothermal tracking (Hedgecock and Russell, 1975). Likewise, when crawling in a chemoattractant gradient, *C. elegans* exhibit gradual steering to orient in the preferred direction, a behavior called klinotaxis, as long as they are at an angle relative to the gradient that allows sensory sampling across head sweeps (Iino and Yoshida, 2009; Kato et al., 2014). Steering and the biased random walk operate on fundamentally different principles (Lockery, 2011). While the latter operates primarily by sensory integration over tens of seconds (Luo et al., 2014a, 2014b; Pierce-Shimomura et al., 1999), steering requires constant, ongoing integration of sensory and motor information on a short time scale (Iino and Yoshida, 2009; Kato et al., 2014).

We previously showed that each of a pair unipolar interneurons called RIA can simultaneously encode both head movements and sensory inputs through two types of calcium signal (Hendricks et al., 2012). Local, compartmentalized calcium events in the ventral and dorsal segments of the RIA axon (nrV and nrD) were correlated with dorsal and ventral head bends, respectively (Figure 1A-B). These axonal compartments correspond to sites of synaptic input from a pair of head motor neurons, the SMDs (White and Southgate, 1986). We will refer to these compartmentalized events within the nerve ring as mCa^2+^, for motor-evoked calcium events. In addition, removal of an attractive odor resulted in calcium increase throughout the entire axon, including nrV and nrD, while odor presentation resulted in whole-axon suppression of calcium levels (Figure 1C). These events will be referred to as sCa^2+^, for sensory-evoked calcium events. These responses are dependent on synaptic input from upstream sensory neurons and interneurons, which synapse onto RIA exclusively in the “loop” region of the axon in the ventral nerve cord (Figure 1A).

**Figure 1.**
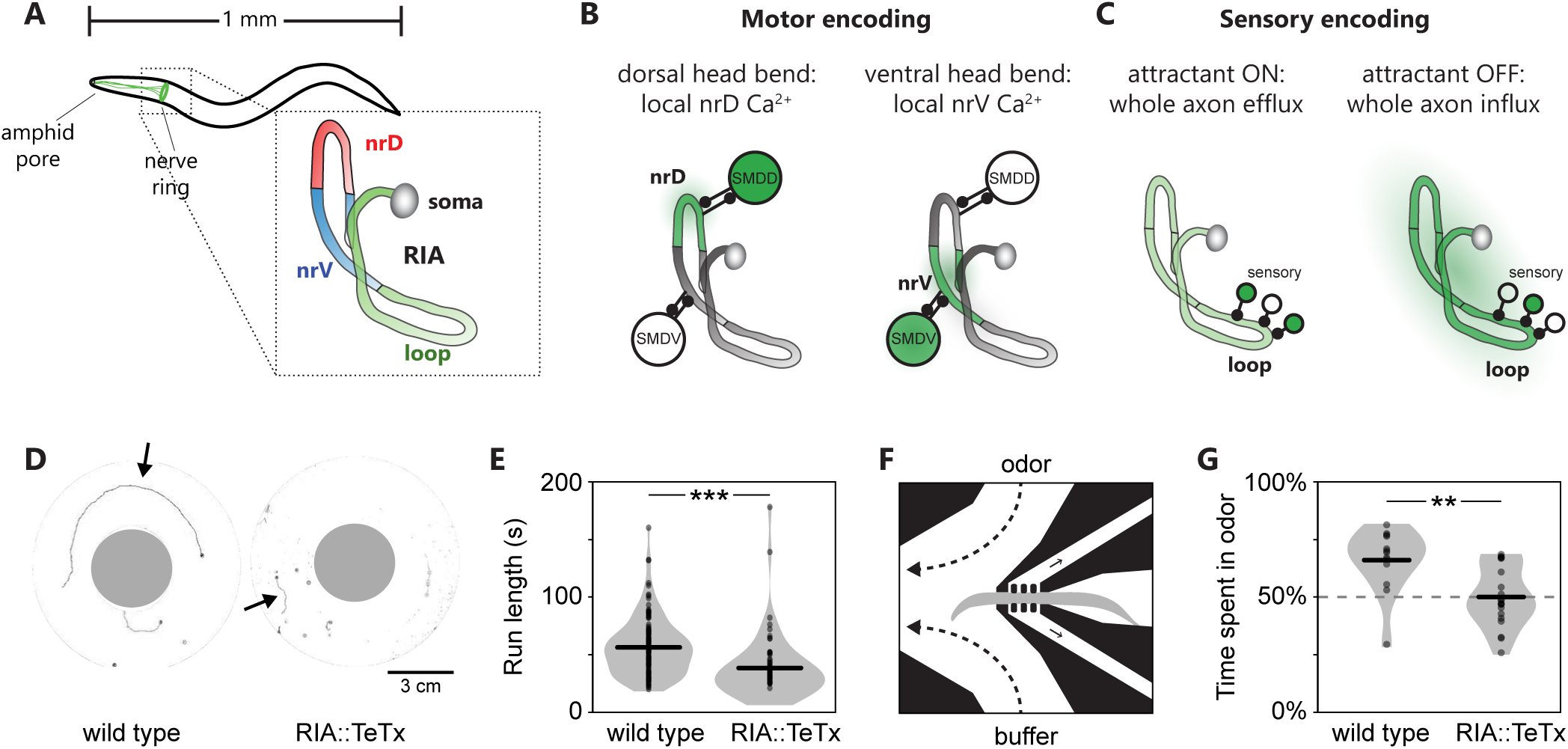
RIA overview and requirement for steering behaviors. (A) RIA is a unipolar interneuron whose axon extends into the ventral nerve cord and throughout the nerve ring. There are two RIAs, left and right, one is shown. (B) Two axonal compartments within the nerve ring, nrV and nrD, encode head movements via local calcium signals (mCa^2+^) triggered by muscarinic input from SMD head motor neurons. (C) Sensory pathways synapse on the loop domain of the RIA axon in the ventral nerve cord. Attractive stimuli lead to whole-axon reduction in calcium, while attractant removal causes whole axon calcium increases (sCa^2+^). (D) Example wild type and RIA::TeTx trajectories over 36 seconds in a radial temperature gradient. (E) Animals expressing TeTx in RIA are defective in isothermal tracking as measured by shorter forward runs (n = 122 wt, 58 RIA::TeTx, t test P < 0.0001). (F) Schematic of head orientation measurement in response to a unilaterally presented stimulus. (G) Wild type animals show a clear preference for odor as measured by percent of the assay time (120s) with their head deflected toward the odor stream, while RIA::TeTx animals do not. n = 13 wt, 20 RIA::TeTx. t-test P = 0.0012.

Within the nerve ring, mCa^2+^ and sCa^2+^ signals overlap spatially and temporally. We showed that these signals were genetically separable, and they were additive with one another within each nerve ring compartment. This additivity suggests a simple mechanism for integrating head orientation and sensory input to calculate the spatial distribution of a stimulus across each head sweep. We therefore proposed a model for steering based on asymmetric output from RIA to head motor neurons (Hendricks and Zhang, 2013).

Here, we confirm and extend aspects of this model and by examining simple head orientation behaviors and sensory encoding events in RIA, which have not been explored in detail and which reveal new features of sensorimotor integration in RIA. We propose a “gate and switch” model for RIA function in which sensory filtering prevents inappropriate activation of RIA when the head is not bent (gating), while mCa^2+^ asymmetry constitutes a switch that routes an inhibitory signal to dorsal or ventral motor neurons to bias gait. Finally, we identify explanatory shortcomings of the model that require further study.

## Results

### RIA in steering and head orientation

Ablating RIA does not grossly affect random walk dynamics in isotropic environments or in response to olfactory stimuli (Gray et al., 2005; Ha et al., 2010). There are seemingly conflicting studies of the role of RIA in response to temperature. One of RIA’s first identified functions was in thermotaxis, as laser ablation led to defective navigation in temperature gradients (Mori and Ohshima, 1995). Later work, also based on laser ablation, found no significant thermotaxis defects (Luo et al., 2014a). However, the assays have a key difference: the former experiments by Mori and Ohsima were conducted in radial temperature gradients, where isothermal tracking dominates thermotactic behavior, while the latter assessed animals in linear temperature gradients, where the biased random walk predominates. These results can be reconciled if RIA is specifically required for steering and not for the biased random walk.

We first confirmed Mori and Ohshima’s isothermal tracking result. Rather than using laser ablation, we expressed the tetanus toxin light chain (TeTx) under the RIA-specific promoter, Pglr-3. TeTx blocks synaptic release by cleaving synaptobrevin (Schiavo et al., 1992). During isothermal tracking, reversals are suppressed and forward runs are extended much longer than during biased random walk-based navigation. This allows run length to be used as a simple metric to measure isothermal tracking behavior (Luo et al., 2007; Ryu and Samuel, 2002). We placed wild type and RIA::TeTx animals in radial temperature gradients (Figure 1D) and recorded their locomotion. Consistent with RIA ablation observations, RIA::TeTx animals did not exhibit gross locomotor defects in isotropic environments, but exhibited a range of defects in a radial temperature gradient, including failure to track, shorter bouts of forward locomotion, and frequent pausing. We focused on run length as a straightforward indicator of isothermal tracking performance. Compared to wild type animals, RIA::TeTx animals exhibited shorter runs, a hallmark of tracking defects (Figure 1E).

Steering behavior depends on asymmetric head bends that propagate through the normal locomotor body wave, producing curved forward movement (Iino and Yoshida, 2009; Izquierdo and Lockery, 2010; Lockery, 2011). To test the idea that RIA is involved in biasing head movements in response to an asymmetrically presented olfactory stimulus, we analysed head movements in a microfluidic chip designed for this purpose (Figure 1F) (McCormick et al., 2011). Because our previous characterization of RIA physiology used the chemoattractant isoamyl alcohol (IAA, 3-Methylbutan-1-ol) and steering has been demonstrated in IAA gradients, we used it as an olfactory stimulus (Hendricks et al., 2012; Kato et al., 2014; Liu et al., 2018). We recorded each animal for two minutes. To control for any intrinsic bias or preference in head bending direction, the odor and buffer streams were switched after one minute. Wild type animals showed a preference for the IAA stream, as measured by the proportion of time spent with their heads bent toward the odor. In contrast, RIA::TeTx animals exhibited no preference (Figure 1G). RIA-defective or ablated animals exhibit normal sCa^2+^ responses and are capable of olfactory chemotaxis, and thus are not sensory impaired (Ha et al., 2010; Hendricks et al., 2012). This is consistent with the hypothesis that RIA is involved in responses to asymmetrically presented stimuli but not the temporal integration the underlies the biased random walk strategy.

### RIA mediates directional head withdrawal

We next wanted to better understand the lower-level behavioral elements that contribute to steering. Specifically, steering requires posture-dependent responses to sensory inputs. We therefore examined the position and velocity of the head immediately before and after IAA removal in semi-restrained animals. While analyzing head movements in restrained animals limits the ability to measure navigation directly, it has the advantage of allowing us to unambiguously assign a stimulus change to a specific point in the head bending cycle. To better visualize oscillatory head bending, we plotted head movements on a 2D position-velocity space that captures its phasic properties (Figure 2A). We then analyzed head movements that occurred immediately after odor removal in relation to each phase of the head bending cycle (Figure 2B). Large ventral head movements were associated with odor removal during dorsal head bends, and odor removal during ventral head bends triggered rapid dorsal head movements (Figure 2C).

**Figure 2.**
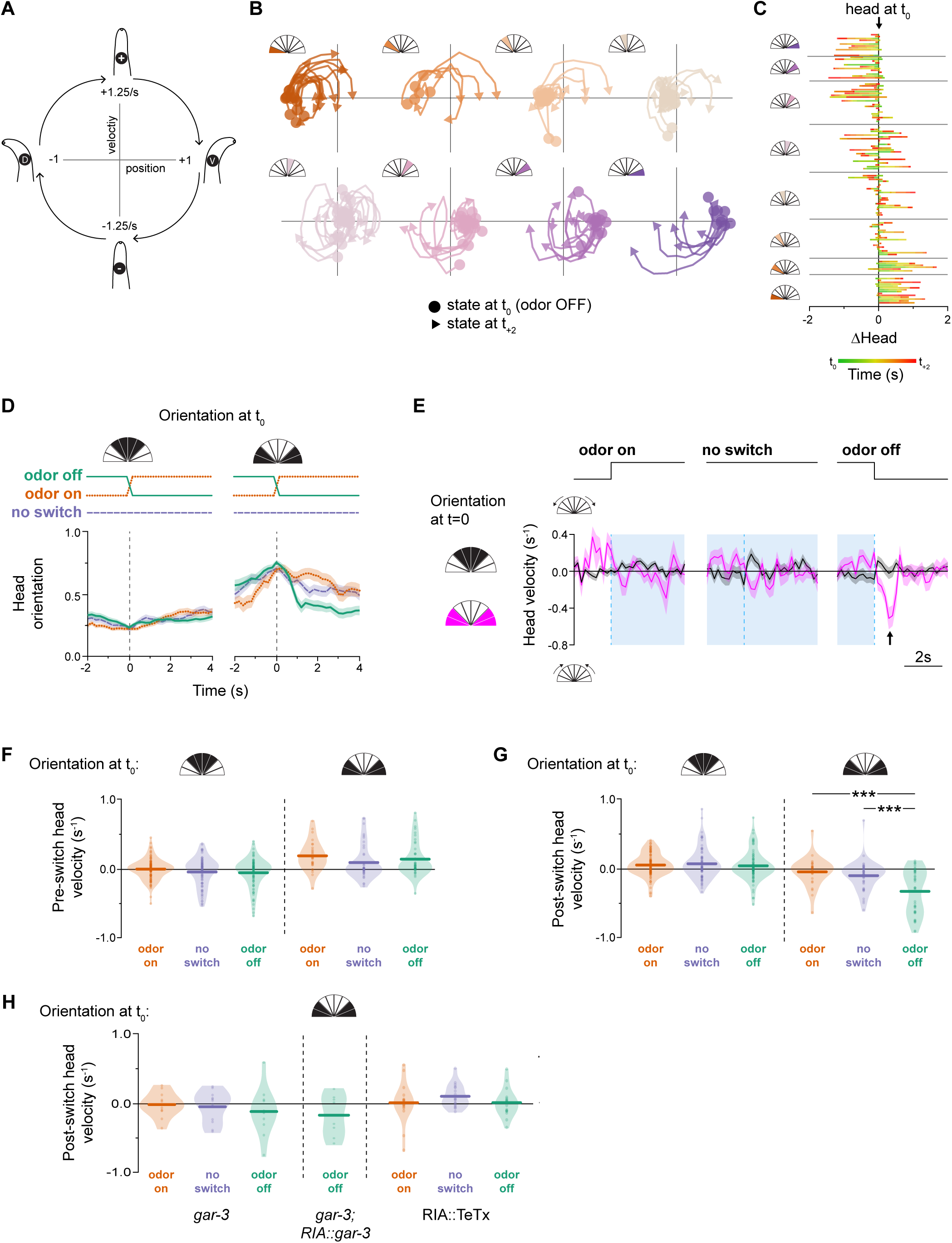
RIA mediates directional head withdrawal. (A) Schematic of position-velocity plots to illustrate oscillatory head movements. (B) Position-velocity trajectories from odor removal (circles) to 2 seconds after (arrowheads). (C) Head displacement in the 2 seconds immediately following odor removal, with the start position normalized to head orientation at the odor switch. (D) Mean plots of peristimulus head movements at odor off, odor on, or in constant odor (no switch), binned according to whether the head is bent (right) or unbent (left) at the time of stimulus change. (E) Comparison of head velocity in response to stimulus changes (or mock switch) when the head is bent or unbent. Positive velocity indicates head deflection, negative velocity indicates head withdrawal. Arrowhead indicates characteristic head with-drawal in response to odor removal when the head is bent. Shading indicates odor presentation. (F) Pre-switch head velocities under each switch and head orientation condition. (G) Post-switch head velocities under each switch and head orientation condition. ANOVA F2,109 = 12.57, P < 0.0001, post-hoc Tukey HSD, odor OFF is significantly different from odor ON (P < 0.0001) and from no switch (P = 0.0002). (H) No head withdrawal response is seen in response to any stimulus switch in *gar-3* mutants (n = 32, ANOVA F2,33 = 0.51, P = 0.6056) or in RIA::TeTx animals (n = 36, ANOVA F2,48 = 0.97, P = 0.3880). Violin plot shading indicates distribution density, bars are means.

Head withdrawal (returning toward center) is expected to follow head bends during oscillatory head movements. To distinguish withdrawal from normal behavioral progression, we compared head movements before and after odor withdrawal to head movements associated with either odor presentation or at an arbitrary time point with no stimulus change. While head bending does tend to be followed by head withdrawal (Figure 2D), as expected, only odor removal elicits a characteristic, transient, rapid head movement and only when the head is bent (Figure 2E,G). Furthermore, rapid head withdrawal was triggered by odor removal irrespective of the direction of head movement just prior to odor removal (Figure 2F). We also examined stimulus-evoked head withdrawal in *gar-3* mutants, which lack mCa^2+^ calcium events, and in RIA::TeTx animals. Neither of these strains exhibited head withdrawals (Figure 2H). However, re-expression of *gar-3* specifically in RIA was not sufficient to restore head withdrawals (discussed below).

### RIA sensory responses are gated according to head position

Models of steering rely on symmetry breaking by phasic modulation of the effects of sensory input on motor networks (Lockery, 2011). Support for a role for RIA in this process was provided in a recent paper in which asymmetric inactivation of RIA synapses produced curved locomotion in the predicted direction (Liu et al., 2018). This relies on generating deeper head bends in the direction of curvature, essentially the inverse of head withdrawal. Both mechanisms rest on the idea that differential mCa^2+^ levels in nrV and nrD to function as a switch to route output from RIA to ventral or dorsal motor neurons. This output is predicted to be inhibitory, and to limit head bends in the unfavorable direction. However restricting head bends must be balanced against the need to maintain forward locomotion. Because mCa^2+^ levels are only partially correlated with head movements and frequently do not show dorsal-ventral differences (Hendricks et al., 2012), there is a potential for inappropriate symmetrical activation or inhibition, which would be predicted to disrupt the animal’s gait.

We therefore analyzed the relationship between head movements, mCa^2+^ asymmetry in the nerve ring, and sCa^2+^ responses in the loop domain. Consistent with previous observations, mCa^2+^ asymmetry (∂nr) shows a clear relationship to head oscillations (Figure 3A). There is no apparent overall relationship between loop sCa^2+^ and head movement (Figure 3B). However, because odor switches occur at set time points in our assay while head movement are spon-taneous, the relationship between head position and odor removal is arbitrary. Spontaneous sCa^2+^ may mask a relationship between head movement and stimulus-evoked events because the latter are much less common. We therefore examined whether stimulus-evoked loop sCa^2+^ events that occur upon odor removal are dependent on head phase. These events trigger calcium influx throughout the nerve ring and are additive with local mCa^2+^ events in nrV and nrD compartments (Hendricks et al., 2012).

**Figure 3.**
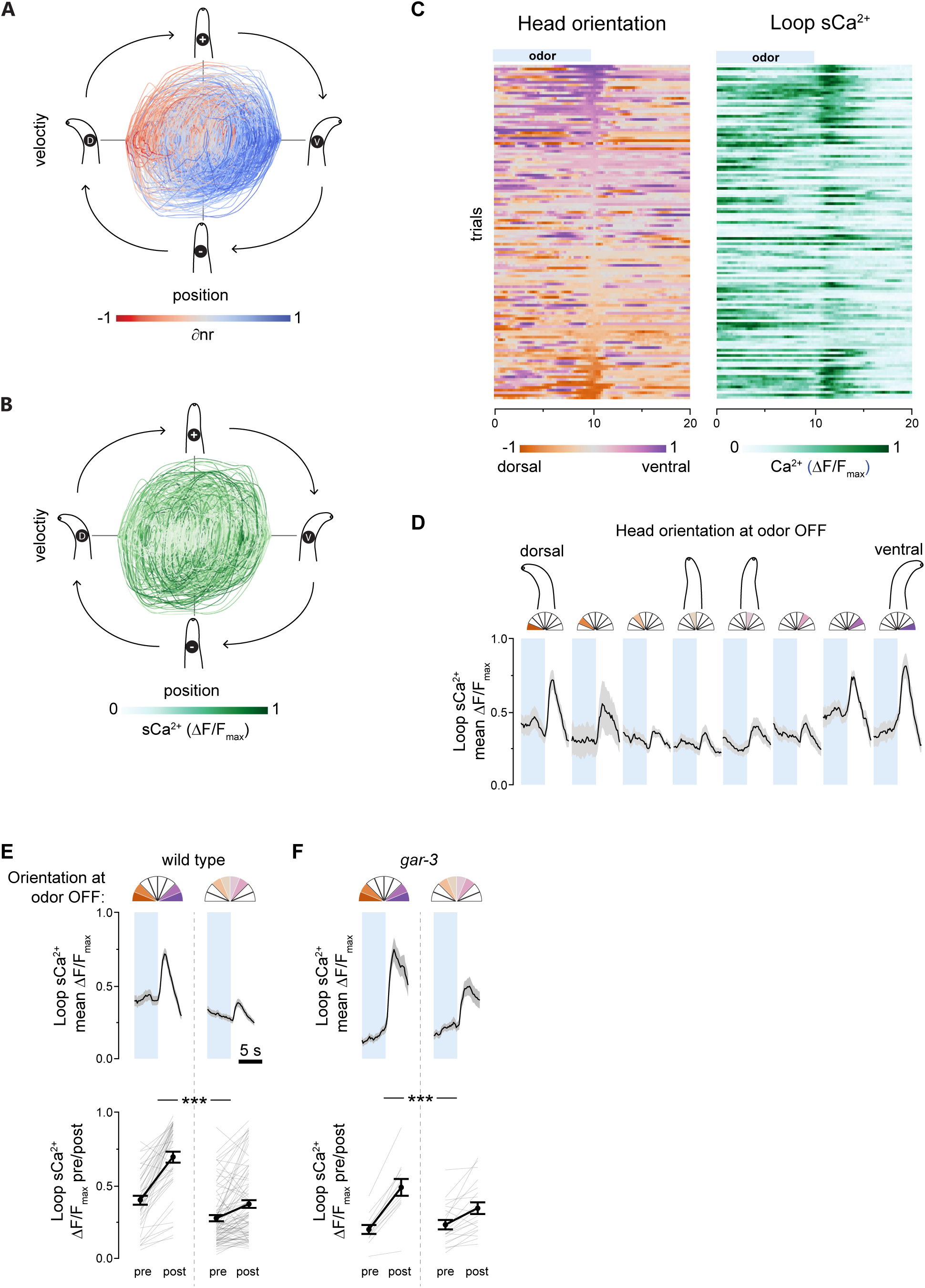
Head orientation gates sensory responses. (A) Head position-velocity trajectories color-coded by ∂nr, a measure of mCa^2+^ asymmetry in nrV and nrD. (B) Head position-velocity trajectories color-coded by loop calcium signal. n = 126, 40 seconds each. (C) Spontaneous head movements (left) and sCa^2+^ signals in the loop region of the RIA axon (right). Rows are matched and sorted according to head orientation at odor off (t = 10s). A linear regression of loop response magnitudes and head deflection in either direction was significant (F1,124 = 29.98, P < 0.0001, n = 126). (D) Mean traces of loop calcium responses 5s before and after odor removal binned by head orientation at the odor off time point. Shading is s.e.m. Left to right, n = 14, 6, 18, 22, 23, 21, 13, 9. (E) Comparison the magnitude of loop calcium responses to odor removal when the head is bent (head orientation > 0.5 or < −0.5) or unbent (head orientation between −0.5 and 0.5). Pre and post measurements are mean loop Ca^2+^ levels in 1s windows just prior to odor OFF and centered on the peak of the mean response, respectively. The response magnitude is larger when the head is bent. N = 42 (bent), 84 (unbent). Repeated measures ANOVA F1,124 = 41.75, P <0.0001. (F) Identical analysis in *gar-3* mutants, which lack compartmentalized nerve ring calcium dynamics. Response magnitudes still depend on head orientation at odor off. n = 12 (bent), 20 (unbent). Repeated measures ANOVA F1,30 = 15.89, P = 0.0004. Blue shading is odor.

We found an unexpected relationship between loop responses and head position at the time of odor removal. Odor removal triggers calcium influx in RIA via inputs from upstream interneurons, and the magnitude of these responses was large when the animal’s head was bent—in either direction— and small or absent when it was not (Figure 3C,D). This affect is the same for dorsal or ventral bends, so for subsequent analyses we defined all bends as positive and classified head positions as “bent” or “not bent” if they were more or less than half the maximal degree of head deflection, respectively. This posture classification revealed a robust difference in the magnitude of sCa^2+^ responses in response to odor removal (Figure 3E).

Posture dependence in behavioral responses to pulsed thermal stimuli was observed by Stephens et al. (2008). In that study, animal posture was defined through a dimensional reduction procedure, and an animal’s state defined by the first two components was predictive of responses to a sudden temperature increase. With only a behavioral readout, it is difficult to tell if posture dependence is based on an active neural mechanism or is a constraint of the motor program. In the case of RIA, we have clear evidence of a sensory gating mechanism. This has important implications for sensory coding in RIA. First, gating allows preferential attention to stimuli encountered during head bends, consistent with a function in steering in response to asymmetrically distributed stimuli. Second, it resolves the issue raised above of potential gait disruption caused by symmetrical output from RIA to head motor neurons. Gating ensures that large sCa^2+^ events are only likely to occur when mCa^2+^ in the nerve ring is asymmetric, i.e. only when the downstream “switch” is engaged to favor either dorsal or ventral synaptic output.

The sensory gating mechanism may act directly on RIA or be a feature of upstream interneurons. We first tested the hypothesis that the gate and switch mechanisms are in fact the same: motor inputs via the muscarinic acetylcholine receptor (mAchR) *Gar-3* that produce local mCa^2+^ events may also somehow gate sCa^2+^ responses. To do so we repeated our analysis of sensory-evoked sCa^2+^ relative to head orientation in *gar-3* mutants. In this mutant, mCa^2+^ events are absent while sCa^2+^ responses are intact, leading to perfectly symmetrical calcium activity in the nerve ring (Hendricks et al., 2012). *gar-3* mutants exhibited normal sensory gating (Figure 3E). This suggests that an independent gating mechanism exists at the level of the RIA loop or upstream interneurons. There are several candidate mechanoreceptor neurons in the head, and this mechanism may rely on a hitherto unidentified proprioceptive pathway.

### Head bending desynchronizes local calcium responses

How sCa^2+^ and mCa^2+^ interact and influence synaptic release is critical for understanding of RIA function. We previously showed that sCa^2+^ and mCa^2+^ are additive within the nerve ring compartments (Hendricks et al., 2012). However, recent work described an observed suppression of mCa^2+^ under the same IAA stimulation pattern (Liu et al., 2018). Both of these papers classified sensory responses by averaging events that were synchronous in all axonal compartments, and neither took into account the effect of head position on sCa^2+^ responses. Thus, filtering in this way cannot distinguish degree of synchronicity from signal strength. Therefore, we analyzed nrV and nrD responses separately during whole-axon sCa^2+^ responses with respect to each other and head velocity. We found that head bending introduced a notable lag in peak responses within the nerve ring that was dependent on head orientation (Figure 4A, B). The polarity of this lag depended on head position, such that the ipsilateral nerve ring compartment exhibited a shorter rise time and earlier peak with respect to the contralateral compartment, consistent with a simple additive model of mCa^2+^ and sCa^2+^.

**Figure 4.**
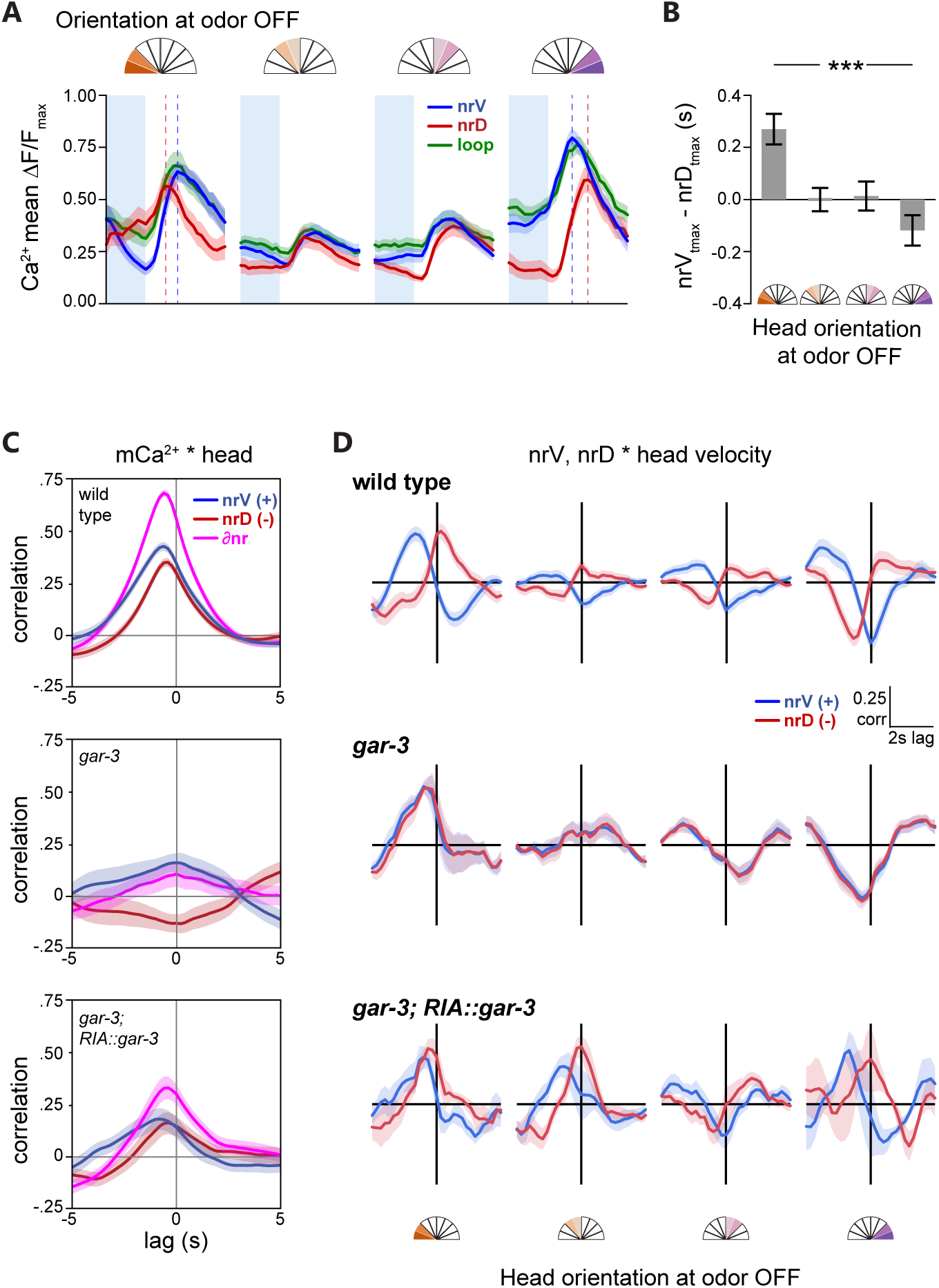
Temporal features of local calcium events in the RIA axon. (A) Mean nrV, nrD, and loop responses at odor removal. Shading is SEM, dashed lines are nrV and nrD mean peaks. (B) Head orientation predicts the mean lag between nrV and nrD peaks post-odor removal. ANOVA F1,124 = 13.47, P = 0.0004. (C) Cross-correlation of nrV, nrD and mCa^2+^ asymmetry (∂nr) and head orientation, showing mCa^2+^ lag with head movements. *gar-3* mutants lack mCa^2+^ and thus do not show head correlations. Re-expression of *gar-3* cDNA in RIA partially rescues this relationship. (D) nrV and nrD cross-correlations with head velocity for wild type, *gar-3*, and RIA-specific *gar-3* rescue in a 6 second time window comprising 2 seconds pre-odor removal and 4 seconds post-odor removal in relation to head orientation at odor off. Peak calcium responses occur with head withdrawal with no lag in the contralateral nerve ring compartment when the head is bent. *gar-3* mutants show no distinction between nerve ring compartments. Expression of *gar-3* cDNA in RIA in *gar-3* mutants does not rescue the temporal features of sensory responses in the nerve ring. Analysis groups are the same as Figure 3.

As previously shown, mCa^2+^ responses typically lag slightly behind head movements (Figure 4C). Possible reasons for this lag include the relative timing of the muscle contractions (via nicotinic acetylcholine receptors) and mCa^2+^ (via muscarinic receptors) elicited by acetylcholine from head motor neurons. To understand how the shift in peak mCa^2+^ related to stimulus-evoked head withdrawal, we performed cross-correlation analysis of head velocities with respect to nrV and nrD calcium dynamics in a restricted time window corresponding to odor removal. During sensory stimulation while the head is bent, peak sensory responses in the nerve ring are coincident with head withdrawal in the contralateral direction but lagged in the ipsilateral direction (Figure 4D). That is, when the head is bent ventrally, peak nrV mCa^2+^ is synchronous with the highest head velocity in the dorsal direction while nrD lags, and vice versa during dorsal head bends. When the head is not bent, this relationship is lost due to the absence of strong sCa^2+^ responses. This desynchronization is lost in *gar-3* mutants, which completely lack mCa^2+^ (Hendricks et al., 2012). Re-expression of *gar-3* specifically in RIA restores some compartmentalization (Figure 4C) but is not sufficient to rescue the temporal structure of stimulus-evoked responses in nrV and nrD, which may relate to its failure to rescue head withdrawal (Figure 4D). This implies either GAR-3 plays additional roles outside RIA in mediating the temporal features of RIA signaling and head withdrawal, or that the rescue construct causes mislocalization, inappropriate expression levels, or otherwise does not sufficiently restore GAR-3 function in RIA. Overall, the additive properties of mCa^2+^ and sCa^2+^ leads to decoupling of local calcium event timing in the nerve ring.

## Discussion

RIA is uniquely situated in the *C. elegans* wiring diagram to integrate head orientation and sensory input. It is one of very few interneurons that receive substantial motor feedback. It receives convergent input from sensory pathways that comprise all *C. elegans* modalities. Here, we confirmed a role for RIA in head orientation in response to an asymmetrically presented stimulus (Liu et al., 2018; Mori and Ohshima, 1995). We identified an elemental component of this behavior, head withdrawal in response to removal of an attractive stimulus. Blocking RIA synaptic release or genetically removing asymmetric local mCa^2+^ in the RIA axon prevented this directional behavior. This is consistent with Liu et al.’s recent demonstration that artificially-induced asymmetric synaptic output from RIA drives steering (Liu et al., 2018). We identified a posture-dependent sensory gating mechanism that may prevent aberrant symmetric RIA output. Phasic behaviors that produce reafferent stimulation often exhibit sensory gating coupled to the behavior. In mice, examples include olfaction, where cholinergic inputs to the olfactory bulb modulate sensory gain in phase with respiration during sniffing, and a functionally similar gating mechanism is evident during whisking (Eggermann et al., 2014; Rothermel et al., 2014). RIA may respond to a similar circuit mechanism that promotes selective attention to relevant stimuli.

The rapid head withdrawal movements we observed, which often involve sudden reversals of head velocity, probably cannot be explained by inhibition of ipsilateral motor neurons alone, but must also involve contralateral muscle contraction. In the nerve ring, the RIA axon receives input from ventral head motor neurons only in nrV and dorsal motor neurons only in nRD. However, output synapses from RIA to SMD and RMD head motor neurons are not segregated in the same way. nrV and nrD both contain interspersed output synapses to both dorsal and ventral motor neurons. However, the post-synaptic anatomical arrangement of these synapses suggest that these connections may be of opposite valence (Figure 5A). Ipsilateral connections are in the distal portion of the axon near neuromuscular junctions, while contralateral synapses are clustered near to the motor neuron cell body. This is at least consistent with the possibility of differing effects of output from nrV and nrD. Exploration of this will require identification of postsynaptic receptors and direct measurement of RIA’s effects on its target neurons.

**Figure 5.**
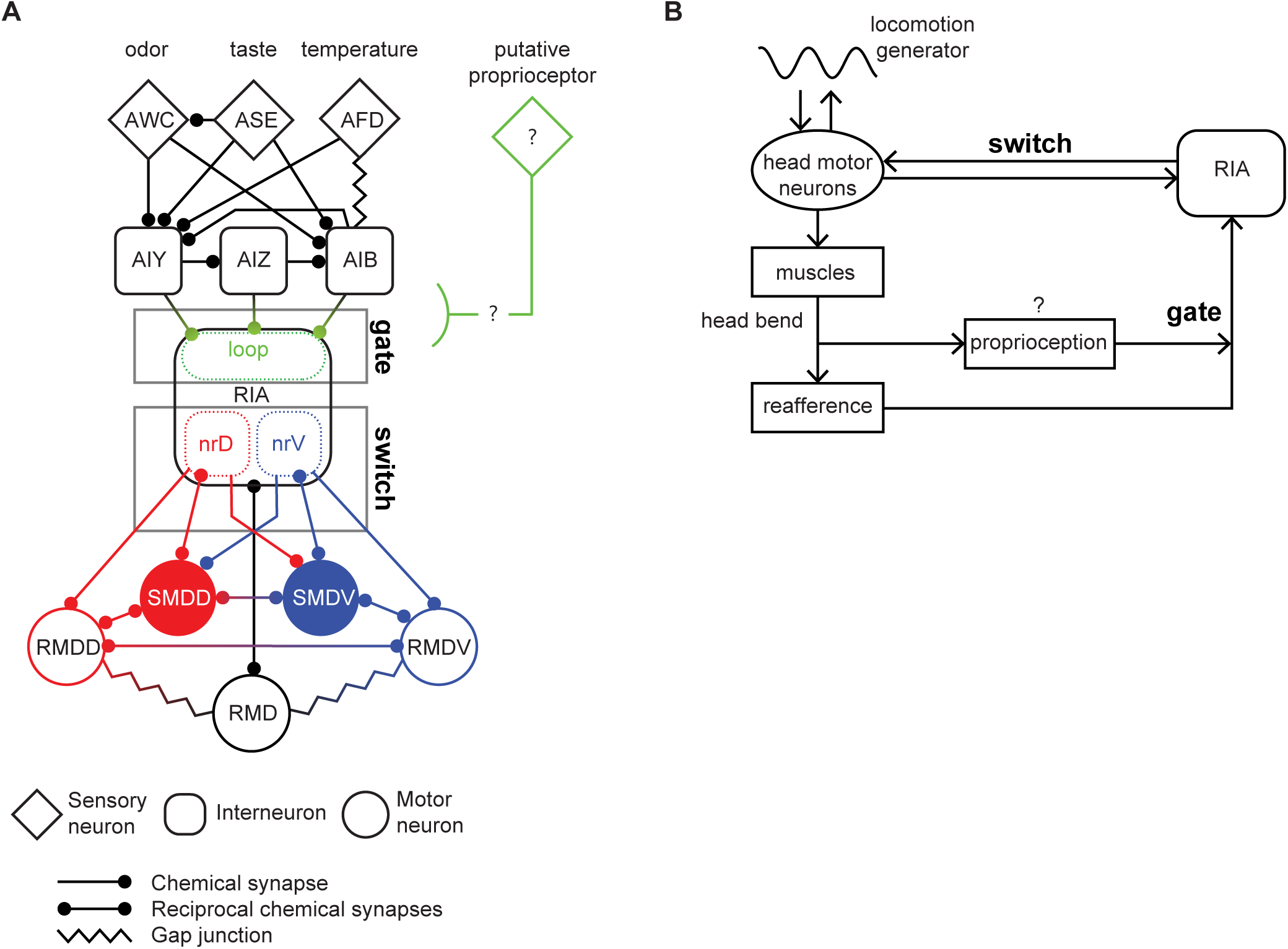
Gate-and-switch model for RIA function in head orientation. (A) RIA circuit diagram, with major upstream sensory pathways and head motor connections. Gating of sensory inputs to the loop domain and the connectivity of the nerve ring connections that constitute the dorsal-ventral switch. (B) Ethological model in which head movements simultaneously produce proprio-ception-gated reafference and efference copies that converge on RIA. RIA output to motor neurons biases the undulatory gait dorsally or ventrally when reafferent stimuli are asymmetric across head sweeps.

Our observation of dynamic sensory gating adds a new layer to the RIA sensorimotor circuit, and resolves potential shortcomings of our previous model, in which RIA outputs but not sensory inputs are regulated by head movements. We propose a two-level “gate and switch” model, in which an as-yet unidentified posture-dependent signal gates sensory sCa^2+^ responses while differential mCa^2+^ within the nerve ring functions as a switch to produce asymmetric synaptic output from the nrV and nrD axonal domains (Figure 5A,B). This switch may depend on the temporal decoupling we observed between nrV and nrD during sensory-evoked responses when the head is bent. mCa^2+^ introduces a temporal bias that reduces the rise time to generate an earlier peak response in the compartment contralateral to the direction of head withdrawal. This peak is coincident with head withdrawal, and the duration of the behavior is similar to the lag introduced between the two axonal compartments. Again, how plausible a mechanism this is will depend on analyzing the effects of these calcium events on synaptic release and its effect on postsynaptic motor neurons.

Despite its small size and simple morphology, RIA exhibits striking computational potential as a sensorimotor integrator. Along with clear behavioral readouts, this relative simplicity makes it an attractive system to probe sensorimotor integration at the cellular level, both in terms of molecular mechanisms and circuit logic.

## Materials and methods

*C. elegans* were raised on nematode growth medium and fed with E. coli strain OP50 according to standard methods (Brenner, 1974; Stiernagle, 2006). Experiments were conducted on young (three day old) adult hermaphrodites. N2; VC657: *gar-3*(gk305) V; MMH099: yxIs19 [Pglr-3a::GCaMP3.3, Punc-122::DsRed]; MMH100: yxIs19; yxIs 20 [Pglr-3a::GCaMP3.3; Pglr-3a::TeTx]; MMH113: yxIs20 [Pglr-3::TeTx::mCherry]; MMH115: yxIs19; *gar-3*(gk305) V. Some strains were provided by the Caenorhabditis Genetics Center (CGC), which is funded by NIH Office of Research Infrastructure Programs (P40 OD010440).

### Isothermal tracking assays

Assay plates were made from fresh, daily-made, thermotaxis agar (2% agar, 0.3% NaCl and 25 mM potassium phosphate buffer pH 6.0) (Goodman et al., 2014). A 2.5 cm glass vial full of frozen glacial acetic acid was placed on the center of each lidless, upside down 9 cm thermotaxis plate 30 minutes prior to the assay to establish a temperature gradient. Worms were grown on standard NGM plates until young adulthood. They were then washed in NGM buffer (1 mM CaCl2, 1 mM MgSO4, 25 mM KPO4 pH 6.0) and placed 1 cm from the edge of the assay plate. Excess liquid was removed and animals were left for 15 minutes before a fresh vial was placed at the center of the plate. Video recordings were made for 20 minutes using an illumination ring made from a red LED strip (Super Bright LEDs, NFLS-X3-LC2) for illumination, Aven Mighty Scope (#2700-200) and VirtualDub software (build 1.10.4). Post-acquisition processing and duration of tracks was assessed using Fiji and Particle tracker plugin (Sbalzarini and Koumoutsakos, 2005; Schindelin et al., 2012).

### Microfluidic device fabrication

Standard soft lithography methods were used to fabricate photoresist (SU8) masters for microfluidic devices (San-Miguel and Lu, 2013). Devices were replica mastered in a two part epoxy resin (Smooth Cast 310, Sculpture Supply Canada #796220) according to the manufacturer’s instructions for long term use. Polydimethylsiloxane (PDMS) (Dow Corning Sylgard 184, Ellsworth Adhesives #4019862)

was mixed at 10:1, degassed, poured over masters, degassed again, and cured at 60°C for at least 3 hours. Inlet holes with made with a Milltex 1 mm biopsy punch (Fisher). Chips were cleaned and then bonded to glass coverslips using air plasma generated by a handheld corona treater (Haubert et al., 2006) (Electro-Technic Products, Chicago, IL). Coupling to fluid reservoirs was done by directly inserting PTFE microbore tubing (Cole-Parmer #EW-06417-21) into inlet holes.

### Calcium imaging

Fluorescence time lapse imaging (100ms exposures, 5 frames per second) was performed as described (Hendricks et al., 2012). Briefly, animals semi-restrained in a microfluidic channel (Chronis et al., 2007) were exposed to alternating streams of NGM buffer or a 1:10,000 dilution of IAA in NGM buffer. Fluid flow was controlled with a ValveBank (AutoMate Scientific). GCaMP3.3 (Tian et al., 2009) intensity was measured from axonal compartments using the Fiji and normalized with the formula (F_t_ - F_min_) / (F_max_ - F_min_). Asymmetry between the nerve compartments (∂nr) was defined as (nrV - nrD) / (nrV + nrD). During calcium imaging, animals were free to move the anterior portion of their head, and these movements were capture as well.

### Head orientation assays

Microfluidic chips were based on a design by McCormick et al. (2011) with minor adjustments to allow stimulus switching halfway through the assay. Animals were exposed to streams of NGM buffer and 100 μM IAA (BioShop ISO900) on either side of their heads. Worms were loaded individually into the chip and once in place their movements were recorded for 2 minutes, with a switch between buffer and IAA at the 1 minute mark, using Aven Mighty Scope (#2700-200) and VirtualDub.

### Head orientation measurements

In both calcium imaging and head orientation devices, head bending was measured at each time point as the angle between the tip of the animal’s nose and the midline of its body at the most anterior restrained point using Matlab. Comparison to previously used methods based on the ratio of pixels on either side of the midline (Hendricks et al., 2012) showed that these approaches yield equivalent results.

## Acknowledgements

This work was supported by funding from McGill University to M.H. and M-H.O., the National Science and Engineering Research Council (NSERC) to M.H. (RGPIN/05117-2014), the Canadian Foundation for Innovation (CFI) to M.H. (32581), and the Canada Research Chairs Program to M.H. (950-231541). We thank Xinyu Liu and Xianke Dong (McGill University) for providing photoresist masters for microfluidic devices. We thank all members of the lab for helpful discussions and comments. Some strains were provided by the Caenorhabditis Genetics Center (CGC), which is funded by NIH Office of Research Infrastructure Programs (P40 OD010440).

## Author Contributions

M.H., M-H.O., and M.J.D. designed the experiments. All authors collected and analyzed data. M.H. and M-H.O. wrote the manuscript.

The authors declare no competing interests.

## References

Berg HC (1975) Bacterial behaviour. Nature 254:389–392.

Brenner S (1974) The Genetic of C. elegans. Genetics 77:71–94.

Calhoun AJ, Murthy M (2017) Quantifying behavior to solve sensorimotor transformations: advances from worms and flies. Curr Opin Neurobiol 46:90–98.

Chronis N, Zimmer M, Bargmann CI (2007) Microfluidics for in vivo imaging of neuronal and behavioral activity in C. elegans. Nat Methods 4:727–731.

Crapse TB, Sommer M a. (2008) Corollary discharge across the animal kingdom. Nat Rev Neurosci 9:587–600.

Eggermann E, Kremer Y, Crochet S, Petersen CCH (2014) Cholinergic signals in mouse barrel cortex during active whisker sensing. Cell Rep 9:1654–1660.

Goodman MB, Klein M, Lasse S, Luo L, Mori I, Samuel A, Sengupta P, Wang D (2014) Thermotaxis navigation behavior. WormBook 1–10.

Gray JM, Hill JJ, Bargmann CI (2005) A circuit for navigation in C. elegans. Proc Natl Acad Sci U S A 102:3184–3191.

Ha H-I, Hendricks M, Shen Y, Gabel CV, Fang-Yen C, Qin Y, Colón-Ramos D, Shen K, Samuel ADT, Zhang Y (2010) Functional Organization of a Neural Network for Aversive Olfactory Learning in C. elegans. Neuron 68:1173–1186.

Haubert K, Drier T, Beebe D (2006) PDMS bonding by means of a portable, low-cost corona system. Lab Chip 6:1548–1549.

Hedgecock EM, Russell RL (1975) Normal and mutant thermotaxis in the nematode C. elegans. Proc Natl Acad Sci U S A 72:4061–4065.

Hendricks M, Ha H, Maffey N, Zhang Y (2012) Compartmentalized calcium dynamics in a C. elegans interneuron encode head movement. Nature 487:99–103.

Hendricks M, Zhang Y (2013) Complex RIA calcium dynamics and its function in navigational behavior. Worm 2:e25546.

Iino Y, Yoshida K (2009) Parallel use of two behavioral mechanisms for chemotaxis in C. elegans. J Neurosci 29:5370–5380.

Izquierdo EJ, Lockery SR (2010) Evolution and analysis of minimal neural circuits for klinotaxis in C. elegans. J Neurosci 30:12908–12917.

Kato S, Xu Y, Cho CE, Abbott LF, Bargmann CI (2014) Temporal responses of C. elegans chemosensory neurons are preserved in behavioral dynamics. Neuron 81:616–628.

Liu H, Yang W, Wu T, Duan F, Soucy E, Jin X, Zhang Y (2018) Cholinergic Sensorimotor Integration Regulates Olfactory Steering. Neuron 97:390–405.e3.

Lockery SR (2011) The computational worm: spatial orientation and its neuronal basis in C. elegans. Curr Opin Neurobiol 21:782–790.

Luo L, Clark DA, Biron D, Mahadevan L, Samuel ADT (2007) Sensorimotor control during isothermal tracking in C. elegans. J Exp Biol 210:3696–3696.

Luo L, Cook N, Venkatachalam V, Martinez-Velazquez LA, Zhang X, Calvo AC, Hawk J, MacInnis BL, Frank M, Ng JHR, Klein M, Gershow M, Hammarlund M, Goodman MB, Colón-Ramos DA, Zhang Y, Samuel ADT (2014a) Bidirectional thermotaxis in C. elegans is mediated by distinct sensorimotor strategies driven by the AFD thermosensory neurons. Proc Natl Acad Sci U S A 111:2776–2781.

Luo L, Wen Q, Ren J, Hendricks M, Gershow M, Qin Y, Greenwood J, Soucy ER, Klein M, Smith-Parker HK, Calvo AC, Colón-Ramos DA, Samuel ADT, Zhang Y (2014b) Dynamic encoding of perception, memory, and movement in a C. elegans chemotaxis circuit. Neuron 82:1115–1128.

McCormick KE, Gaertner BE, Sottile M, Phillips PC, Lockery SR (2011) Microfluidic devices for analysis of spatial orientation behaviors in semi-restrained C. elegans. PLoS One 6:e25710.

Mori I, Ohshima Y (1995) Neural regulation of thermotaxis in C. elegans. Nature 376:344–348.

Pierce-Shimomura JT, Morse TM, Lockery SR (1999) The fundamental role of pirouettes in C. elegans chemotaxis. J Neurosci 19:9557–9569.

Rothermel M, Carey RM, Puche A, Shipley MT, Wachowiak M (2014) Cholinergic inputs from Basal forebrain add an excitatory bias to odor coding in the olfactory bulb. J Neurosci 34:4654–4664.

Ryu WS, Samuel ADT (2002) Thermotaxis in C. elegans Analyzed by Measuring Responses to Defined Thermal Stimuli. J Neurosci 22:5727–5733.

San-Miguel A, Lu H (2013) Microfluidics as a tool for C. elegans research. WormBook 1–19.

Sbalzarini IF, Koumoutsakos P (2005) Feature point tracking and trajectory analysis for video imaging in cell biology. J Struct Biol 151:182–195.

Schiavo G, Benfenati F, Poulain B, Rossetto O, Polverino De Laureto, DasGupta BR, Montecucco C (1992) Tetanus and botulinum-B neurotoxins block neurotransmitter release by proteolytic cleavage of synaptobrevin. Nature 359:832–835.

Schindelin J, Arganda-Carreras I, Frise E, Kaynig V, Longair M, Pietzsch T, Preibisch S, Rueden C, Saalfeld S, Schmid B, Tinevez J-Y, White DJ, Hartenstein V, Eliceiri K, Tomancak P, Cardona A (2012) Fiji: an open-source platform for biological-image analysis. Nat Methods 9:676–682.

Stephens GJ, Johnson-Kerner B, Bialek W, Ryu WS (2008) Dimensionality and dynamics in the behavior of C. elegans. PLoS Comput Biol 4:e1000028.

Stiernagle T (2006) Maintenance of C. elegans. WormBook 1–11.

Straka H, Simmers J, Chagnaud BP (2018) A New Perspective on Predictive Motor Signaling. Curr Biol 28:R232–R243.

Tian L, Hires SA, Mao T, Huber D, Chiappe ME, Chalasani SH, Petreanu L, Akerboom J, McKinney SA, Schreiter ER, Bargmann CI, Jayaraman V, Svoboda K, Looger LL (2009) Imaging neural activity in worms, flies and mice with improved GCaMP calcium indicators. Nat Methods 6:875–881.

White JG, Southgate E (1986) The structure of the nervous system of the nematode C. elegans. Philos Trans R Soc Lond B Biol Sci 314:1–340.

